# Development of the adult-like larval stomach of *Lepidobatrachus laevis*

**DOI:** 10.1101/2022.04.05.487172

**Authors:** Jennifer K. Austiff, James Hanken

## Abstract

Most frogs’ diets shift between the larval and adult phases, from a filter-feeding herbivore to a bulk-feeding carnivore. This change in diet corresponds to a biphasic mode of gut development that begins during embryogenesis and resumes at metamorphosis, when nearly the entire anatomy of the tadpole is reorganized into its adult morphology. The frog *Lepidobatrachus laevis* forgoes this metamorphic shift in feeding mode and instead consumes a bulk-feeding, carnivorous diet both as a larvae and as an adult. This unusual larval diet is enabled by the presence of an adult-like stomach in the tadpole. This study investigates the anatomy and embryonic development of the larval stomach of *L. laevis* and what, if any, further changes occur during metamorphosis. The histology of embryonic and metamorphic stomach development is compared to that of *Xenopus tropicalis*, a frog with a typical larval stomach. We find that *L. laevis* directly forms an adult-like stomach during embryonic development without first forming a larval-like configuration. Moreover, no additional major remodeling of the stomach occurs during metamorphosis, although the stomach does gradually and slightly increase in complexity, proliferating more glands and increasing connective tissue and muscle layers, between hatching and the end of metamorphosis. This developmental trajectory of the stomach in *L. laevis* corresponds with the megalophagous, carnivorous diet these frogs maintain from tadpole to adult, as well as the maintenance of active feeding throughout metamorphosis. These results will facilitate future investigations into the mechanisms underlying the evolution of this unusual larval anuran feeding strategy, as well as the broader study of how development mediates evolutionary change.

## Introduction

The evolution of a complex life history that comprises distinct larval and adult ontogenetic stages is an important ecological strategy. Complex life histories allow discrete life stages of the same species to exploit different niches and reduce competition between offspring and adults (Ziermann *et al*., 2013; Schalk *et al*., 2014). Distinct larval and adult niches, along with the associated morphological and physiological traits, are an adaptation to different selective pressures acting across changing body size. For example, this allows larvae to exploit the seasonal abundance of aquatic primary productivity to facilitate early growth (Wassersug, 1975; Wilbur, 1980; Werner & Gilliam, 1984; Werner, 1986). Divergent larval and adult phases may also enable independent evolution of larval and adult developmental modules, which experience different selective pressures (Hanken, 1993). Such is the case for many frogs. Tadpoles of most species are aquatic, herbivorous/omnivorous, microphagous feeders (i.e., they are filter-feeders or otherwise ingest very small food particles) while adult frogs typically are terrestrial, carnivorous, macrophagous feeders (i.e., they consume bulky food items) (Ruibal & Thomas, 1988). This developmental trajectory is highly conserved among anuran species and is the plesiomorphic state for the group (Griffiths, 1961). The shift in diet associated with metamorphosis from larva to adult corresponds to significant transformation of many aspects of frog morphology, including the cranial skeleton and musculature, oral apparatus and gastrointestinal tract (Ziermann *et al*., 2013; Schalk *et al*., 2014).

The stomach is of particular interest, as most species of metamorphosing frogs exhibit a biphasic pattern of gut development that is correlated with the shift in diet from larva to adult. Vertebrate stomachs typically consist of an anterior glandular region, the fundic stomach, that secretes digestive enzymes and stomach acid, and a posterior muscular region, the pyloric stomach, that mechanically churns the food and enzymes (Andrew & Hickman, 1974; Shi & Ishizuya-Oka, 1996; Schreiber *et al*., 2009). Indeed, this configuration is observed in adult frogs, which typically are active carnivores that favor large animal prey, and where functionally specialized fundic and pyloric regions aid the digestion of bulky food items (Shi & Ishizuya-Oka, 1996; Smith *et al*., 2000; Schreiber *et al*., 2009). However, tadpoles of most anuran species instead have a reduced stomach with a glandular structure, or *“manicotto glandulare”* (Lambertini, 1946; Griffiths, 1961). The larval foregut also has a slightly basic pH (7.8) and lacks any peptic, amylase or lipase activity (Barrington, 1945; Lambertini, 1946; Dodd & Dodd, 1976; Fox, 1983; Griffiths, 1961; Rovira, *et al*., 1993). This stomach morphology is functionally well suited for digesting the microphagous, herbivorous/omnivorous diet that is characteristic of the tadpoles of most species of frogs. For microphagous feeding, there is no need for a large, enzymatically active stomach that breaks down food. Instead, the intestine is greatly elongated, providing increased surface area for nutrient absorption of small particles (Barrington, 1942, 1945; Fox, 1983). The transition from larval to adult stomach occurs during metamorphosis, when the lining of the larval gastrointestinal tract sloughs off and the adult lining forms anew from stem cells deep within the gut wall (Shi & Ishizuya-Oka, 1996; Schreiber *et al*., 2009). These stem cells give rise to a new stomach lining with functionally and morphologically distinct fundic and pyloric regions. Not surprisingly, feeding does not occur during this transition but instead resumes once adult stomach components are in place and functional.

Although this ontogenetic trajectory is widely conserved among anurans, larval development and ecology may vary significantly, and this variation can have profound impacts on stomach development and morphology (Womble, *et al*., 2016). Direct-developing frogs, for example, bypass the free-living, aquatic larval phase and instead feed initially only on abundant yolk reserves within the egg, hatching into miniature adults. This developmental strategy has evolved many times in anurans (Grosjean, *et al*., 2008). At least some direct-developing species never form a functional larval foregut. Instead, they form an adult digestive tract during embryogenesis that is derived from a different population of cells than the one that forms the gut in most metamorphosing frogs (Buchholz, *et al*., 2007; Karadge & Elinson, 2013). Spadefoot toad tadpoles, on the other hand, initially form a typical tadpole foregut, but under certain environmental pressures may transform into a carnivorous larval morph with an adult-like stomach and mouth. This transformation facilitates feeding on smaller siblings and other animal prey (Pfennig, 1992).

One of the most extreme departures from the ancestral, biphasic pattern of stomach development characteristic of most metamorphosing frogs is seen in the South American genus *Lepidobatrachus*, which comprises three species endemic to the Chacoan desert of Argentina, Paraguay and Bolivia (Budgett, 1899; Reig & Cei, 1963). Tadpoles of *L. laevis*, the most extensively studied species, hatch with a seemingly adult-like trophic morphology—mouth, jaws, and gut—that enables a macrophagous, carnivorous diet so extreme that it has been termed “megalophagy” (Ruibal & Thomas, 1988). The tadpole is able to capture, ingest and digest live animal prey almost as large as itself (Ruibal & Thomas, 1988; Carroll *et al*., 1991; Fabrezi *et al*., 2016). They also are able to feed throughout metamorphosis (Ruibal & Thomas, 1988; Fabrezi & Cruz 2020). The ability to consume animal prey continuously from hatching through and beyond metamorphosis, paired with previous studies of larval stomach histology (Ruibal & Thomas, 1988), suggests that *L. laevis* does not form a typical larval stomach during embryogenesis, but instead forms an adult-like stomach that does not transform significantly during metamorphosis. These predictions, however, have never been confirmed by direct examination of stomach development across multiple developmental stages. In this study, we ask 1) What is the sequence and relative timing of development of the major stomach tissues in *L. laevis?* Does the stomach develop directly into an adult-like configuration, or are any larval-specific stomach elements recapitulated or even retained? 2) Does the stomach undergo any significant remodeling during metamorphosis, and, if so, in what ways? We describe embryonic stomach development in *L. laevis* and compare it to development of the larval stomach in *Xenopus tropicalis*, a species that exhibits the biphasic pattern of gut development characteristic of most metamorphosing anurans (Sterling *et al*., 2012). Additionally, we compare stomach metamorphosis between *L. laevis* and *X. tropicalis*. Stomach development of both species is characterized at both gross and histological levels. By generating a more precise and detailed developmental timeline for *L. laevis*, we hope to gain greater understanding of its novel gastric ontogeny, as well provide a more extensive baseline for future studies that explore the genetic and developmental mechanisms that underlie the heterochronic shift of a typically adult trait to the larval period.

## Methods

### Breeding

Adult *Xenopus tropicalis* were obtained from the Hanken Lab at Harvard University. Adult frogs were housed individually in reverse-osmosis (RO) water at 28°C. A pair of adults were placed in a 10-gallon tank filled with 10% Holtfreter solution (Holtfreter, 1931) and injected with 10 international units (IU) of human chorionic gonadotropin (HCG; Sigma CG10) into the dorsal lymph sac. The following day, females were injected with 300 IU HCG and males with 100 IU, which induced amplexus. Once eggs were laid, adults were returned to individual tanks and the eggs collected. Fertile eggs were separated into groups of 25 and placed in 30-mm petri dishes in 10% Holtfreter solution. After *NF 45*, tadpoles were raised together in 10% Holtfreter solution in a 10-gallon tank at 28°C. They were fed phytoplankton (Sera Micron) once per day after the onset of feeding.

Adult *Lepidobatrachus laevis* were obtained from the Nascone-Yoder Lab at North Carolina State University, College of Veterinary Medicine. Breeding protocol followed Amin (2015). Adult frogs are housed individually in dechlorinated tap water at 28°C. Adult females were induced to breed by injecting 0.5 mL (50 μg) luteinizing hormone-releasing hormone (LHRH; Sigma L7134) into the dorsal lymph sac. Adult males were injected with 0.3 mL (30 μg) LHRH 1.5 hr later. The male was then introduced into a 10-gallon breeding tank, followed by the female. The tank was covered with a towel, and frogs were checked every hour for signs of aggression or amplexus. Once amplexus was observed and eggs were deposited, adults were returned to individual tanks and the eggs collected. Fertile eggs were separated into groups of 25 and placed in 100-mm petri dishes in 10% Holtfreter solution. After hatching, larvae were separated into pairs and placed in condiment cups incubated at 28°C. Once approximately 1.5 cm in length, tadpoles were raised individually in 1-mL pipette-tip boxes. Between stages *GS 32* and *GS 36*, tadpoles were transferred through decreasing concentrations of Holtfreter solution until they were in pure dechlorinated tap water. The tadpoles were fed a diet of smaller *L. laevis* tadpoles and *X. laevis* tadpoles (Xenopus Express). Just before and during metamorphosis, the diet was supplemented with redworms and nightcrawlers (BassPro Shop).

### Staging

*Xenopus tropicalis* embryos and tadpoles were staged according to the morphological criteria of Nieuwkoop & Faber (NF; 1994), which defines 66 stages between fertilization and the end of metamorphosis. *Lepidobatrachus laevis* specimens were staged according to the morphological criteria of Gosner (GS; 1960), which defines 46 stages over the same interval. The two staging schemes were compared to determine equivalent developmental stages across embryogenesis and metamorphosis. Samples of *X. tropicalis* were collected at NF stages 41, 43, 45, 53, 60, 63 and 65. Samples of *L. laevis* were collected at Gosner stages 21, 23, 25, 31, 41, 44 and 46.

### Fixation and preservation

Embryos and tadpoles were euthanized by immersion in 2% tricaine mesylate (MS-222; Sigma E10521). They were fixed in Bouin solution (Cold Spring Harb Protoc, 2006; Sigma HT10132) for 24 hr as whole specimens or, when tadpoles were larger, as dissected stomachs. Specimens were rinsed in 1X phosphate-buffered saline (PBS; Cold Spring Harb Protoc, 2006) until residual yellow coloration from Bouin solution was eliminated, then dehydrated in 70% ethanol and stored at −20°C.

### Sectioning

Specimens were dehydrated by immersion in 100% ethanol, transferred to xylene, infiltrated with Paraplast (Sigma P3558) using the Tissue Tec, and embedded in Paraplast. Specimens were serially sectioned at 7-μm thickness using a Leitz 1512 microtome, and the sections mounted on glass slides.

### Staining

Mounted sections were cleared with xylene, then rehydrated through an ethanol series. Sections were stained using the Mallory trichrome method (Presnell *et al*., 1997) as modified by Lewis & Hanken (2016). Slides were stained with Mayer hematoxylin for 10 min, washed in running rRO water for 10 min, stained in Mallory I (1% acid fuchsin) for 30 sec, rinsed briefly in RO water, stained in 1% phosphomolybdic acid for 5 min, and then stained in Mallory II (1% orange G, 1% aniline blue, and 2% oxalic acid) for 3 min before a final rinse in RO water. Sections were dehydrated through an ethanol series, cleared with xylene, mounted with EZ-mount mounting medium (Epredia 9999120) and sealed with a coverslip.

### Imaging

Sections were imaged on a Leica DMRE microscope using a Leica DMC5400 camera (Wetzlar, Germany) and LAS X software (RRID: SCR 013673; Leica Microsystem). Images were edited using Fiji for Mac OS X on ImageJ, version 1.51 (Preibisch *et al*., 2009; Scheindlin *et al*., 2012).

## Results

### Development of the larval stomach

#### Xenopus tropicalis

At *NF 41*, the gut is a thick, yolky tube (Fig. 1A). As seen in ventral view, there is a slight indentation in the anterior, left side as the gut bends slightly to the right but then straightens as it descends caudally. This indentation, or gastroduodenal loop (GD), marks the boundary between the foregut and midgut. Further posteriorly, the midgut narrows and transitions into the hindgut. The microanatomy of the foregut, midgut and hindgut are indistinguishable (Fig. 2A–D). The entire gut tube is a nearly solid mass of undifferentiated yolky endodermal cells; the gut lumen and liver diverticulum are just beginning to form.

**Figure 1.**
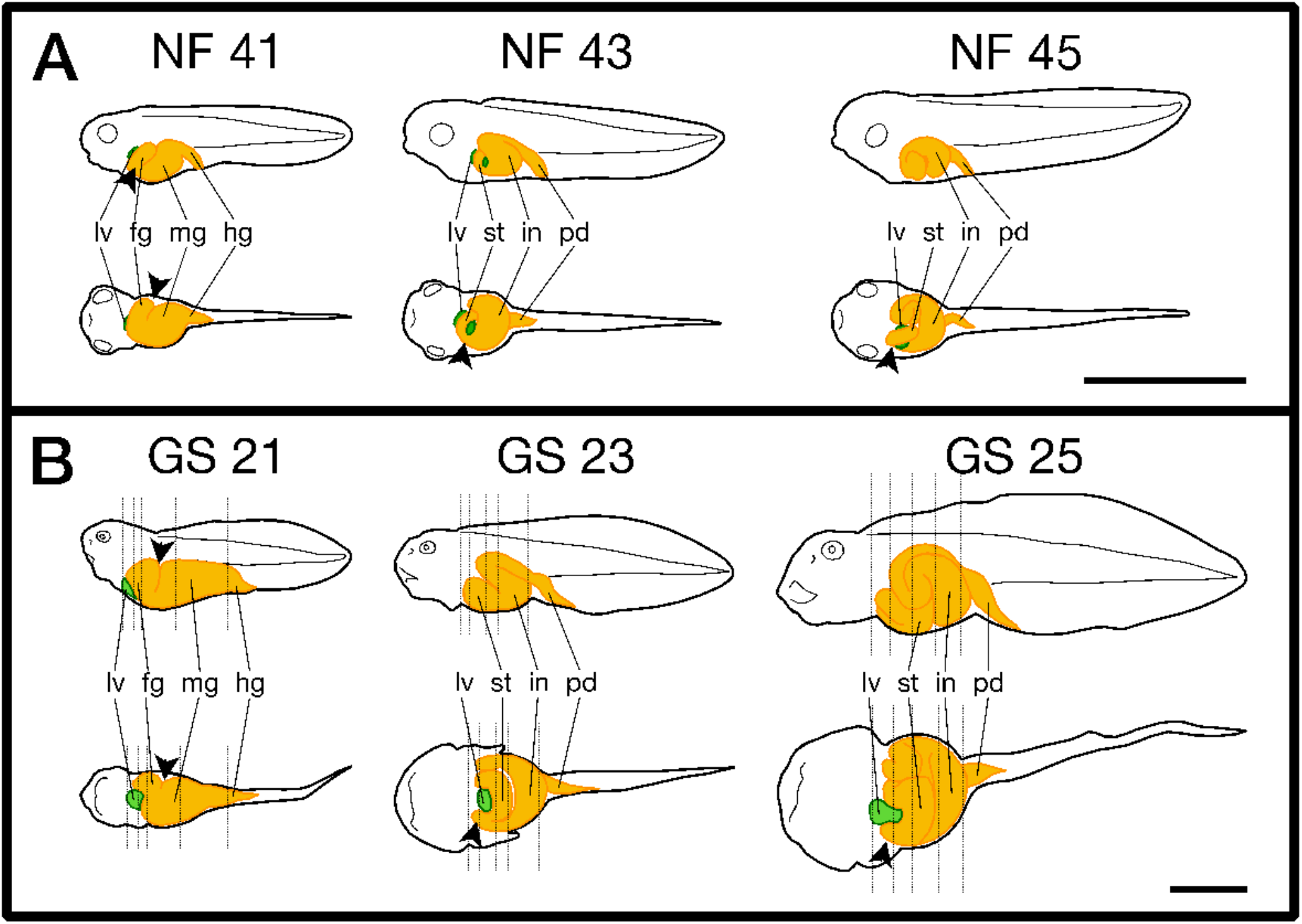
Differences in the relative timing of early gut development between (A) *Xenopus tropicalis* and (B) *Lepidobatrachus laevis*. The gut (yellow) consists of foregut (fg), midgut (mg) and hindgut (hg), or stomach (st), intestine (in) and proctodeum (pd), depending on stage. The gastroduodenal loop, where visible, is indicated by an arrowhead. The liver (lv) is green. In each panel, the top row depicts left lateral views; the bottom row depicts ventral views. Anterior is to the left. With respect to gut development, the three embryonic stages shown for *L. laevis* (Gosner; GS) are approximately equivalent to the three larval (posthatching) stages shown for *X. tropicalis* (Nieuwkoop & Faber; NF). Hatching occurs at *GS 25* and *NF 35*, respectively. Scale bars: A, 2 mm; B, 2 cm.

**Figure 2.**
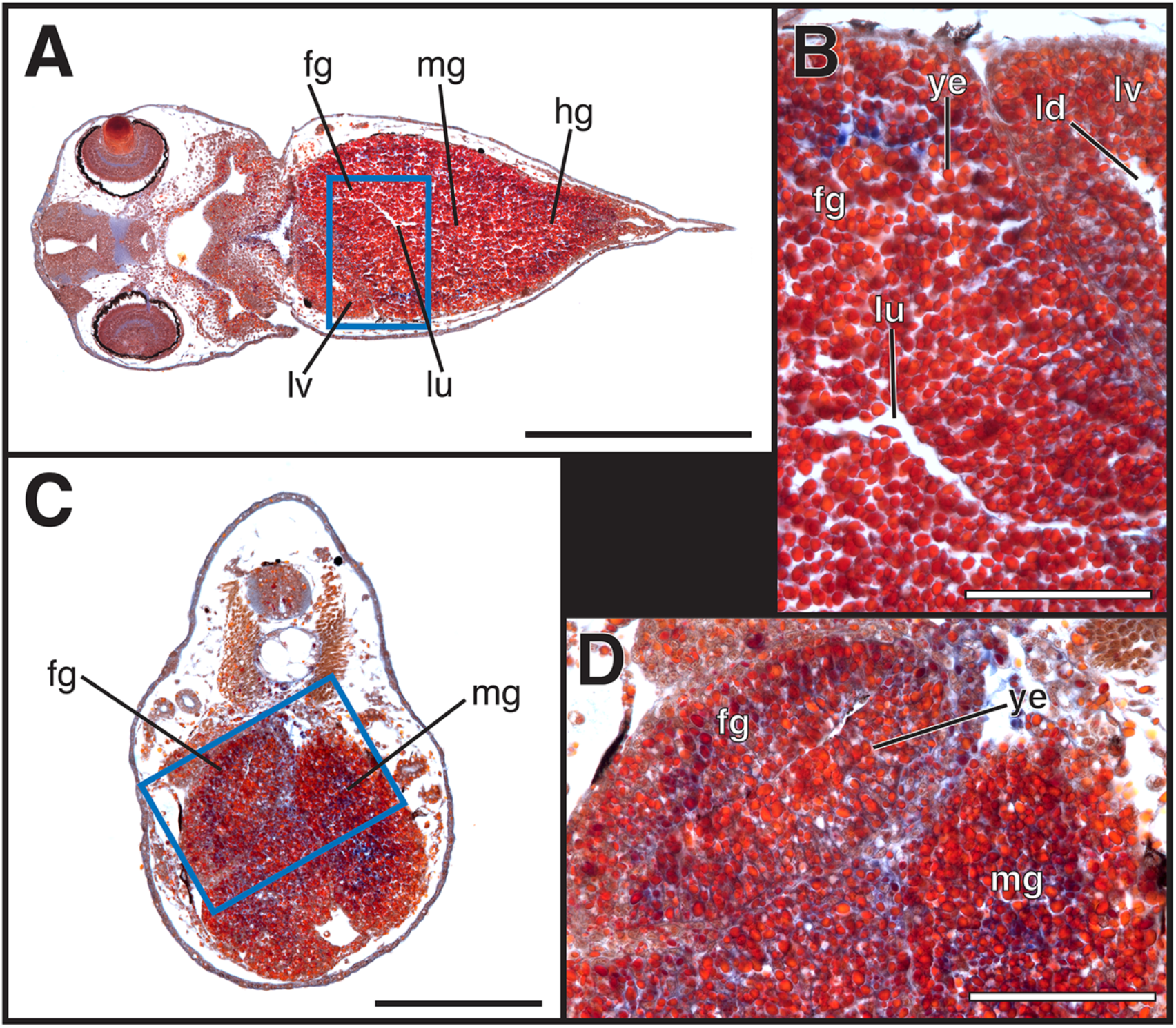
*Xenopus tropicalis* embryo, *NF 41*. A) Coronal section; anterior is to the left. The foregut (fg), recognizable by the incipient lumen (lu), is followed posteriorly by the midgut (mg) and hindgut (hg). The liver (lv) is also visible. Area enclosed by the blue rectangle is enlarged in B. B) The foregut (fg) and liver (lv) consist of yolky endodermal cells (ye) surrounding the lumen (lu) and diverticulum (ld), respectively. C) Transverse section; dorsal is at the top. The foregut (fg) is on the left side of the specimen; the midgut (mg) is on the right side and ventral. Area enclosed by the blue rectangle is enlarged in D. D) The foregut (fg) and midgut (mg) consist of yolky endodermal cells (ye). Magnification: A and C, 20X; C and D, 40X. Scale bars: A, 500 μm; B, 100 μm; C, 200 μm; D, 100 μm. Staining: yolky endoderm, orange to dark red; muscle, red; connective tissues, epithelium and mucous, blue; red blood cells, yellow to orange.

By *NF 43*, the gut has begun to elongate and loop, and it continues to differentiate into discrete organs. The anteriormost portion is the esophagus. Posterior to the esophagus, the gut tube transitions to become the presumptive larval stomach (*manicotto glandulare*), which bends to the embryo’s right side as it loops ventrally around the liver (Fig. 3A). The gut then loops back dorsally, transitioning from stomach to intestine. After this loop, the gut curves transversely to the left and thickens considerably. Then it bends anteriorly as it loops ventrally, ending caudally in the proctodeum (Fig. 1A). The microanatomy of the stomach and of the intestine are beginning to diverge. The stomach has started to organize into a simple columnar epithelium, with ciliated cells facing a central lumen. This epithelium is homogeneous, and most of the cells are still yolk-filled. An outermost serosal layer can also be distinguished (Fig. 3B). The lumen extends into the intestine. There is no distinct epithelium in the intestine, but the yolky endodermal cells have begun to organize into columns (Fig. 3C).

**Figure 3.**
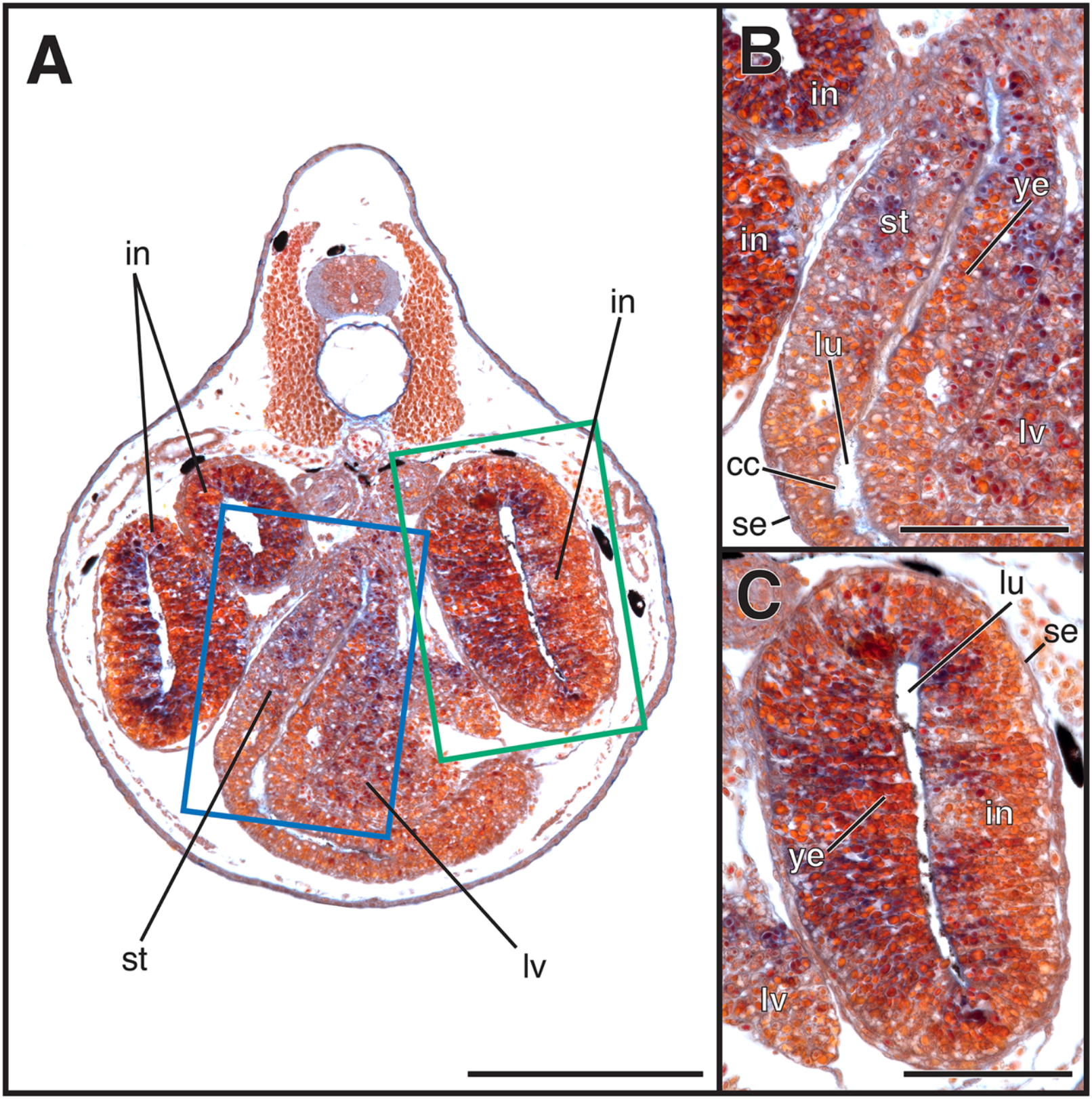
*Xenopus tropicalis* embryo, *NF 43*. A) Transverse section; dorsal is at the top, The presumptive stomach (st) is in the center of the visceral cavity, flanked on the embryo’s right side by one loop of the intestine (in) and on the left side by two loops. The liver (lv) is nestled in a bend of the stomach on the embryo’s right side. Areas enclosed by the blue and green rectangles are enlarged in B and C, respectively. B) Ciliated cells (cc) line the lumen (lu) of the developing stomach (st), but much of the gut tube is still yolky endoderm (ye) surrounded by an outer serosa (se). C) The intestine (in) consists of yolky endoderm (ye) surrounding a central lumen (lu) with external serosa (se). Magnification: A, 20X; B–C, 40X. Scale bars: A, 200 μm; B–C, 100 μm.

At *NF 45*, the gut is a long coiled tube. The stomach is very short; in ventral view, it lies dorsal to the intestine, which spirals one-and-a-half revolutions (Fig. 1A). The intestine is greatly elongated. In transverse section, its extensive looping is visible as many cross sections to the left of and ventral to the stomach; the liver sits adjacent and dorsal to the stomach (Fig. 4A). Major regions of the gut are now clearly differentiated histologically. The stomach has begun to develop gastric glands and a ciliated epithelium (Fig. 4B). There is no muscularis mucosa or submucosa present. The intestine lags behind in development: it is still disorganized with patches of yolky cells, but the epithelium is differentiated (Fig. 4C).

**Figure 4.**
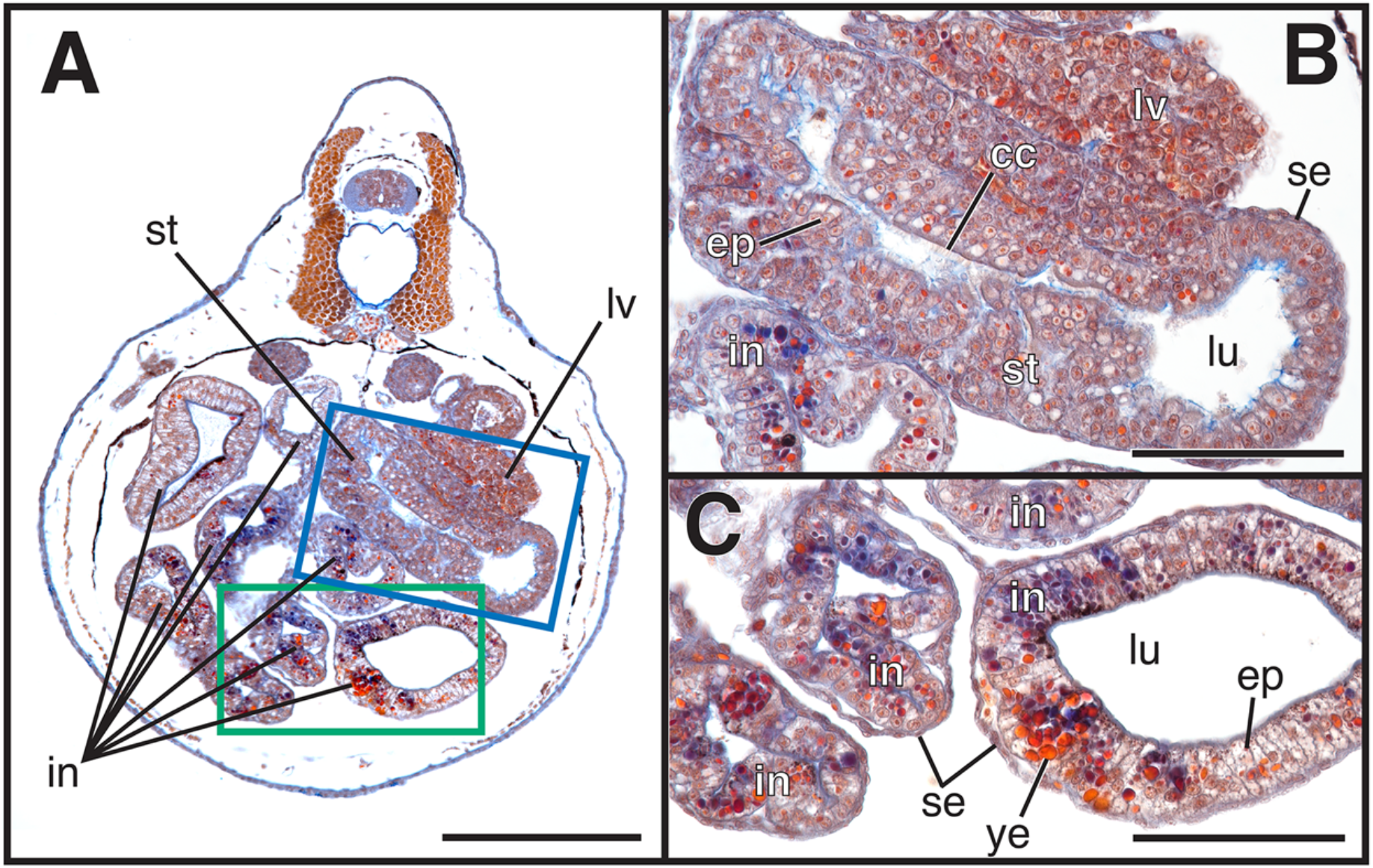
*Xenopus tropicalis* embryo, *NF 45*. A) Transverse section; dorsal is at the top. The stomach (st) dominates the right side of the visceral cavity, where it is flanked dorsally by the liver (lv). Several loops of the elongate intestine (in) are visible to the left of and ventral to the stomach. Areas enclosed by the blue and green rectangles are enlarged in B and C, respectively. B) The stomach is lined internally by a thick, folded epithelium (ep) with ciliated cells (cc) facing the lumen (lu) and is surrounded externally by the serosa (se). The liver (lv) and intestine (in) are adjacent to the stomach. C) The intestine (in) comprises mostly columnar epithelium (ep) facing the lumen (lu) but retains pockets of yolky endodermal cells (ye) surrounded by the serosa (se). Magnification: A, 20X; B–C, 40X. Scale bar: A, 25 μm; B–C, 100 μm.

#### Lepidobatrachus laevis

At *GS 21*, the gut is a thick, yolky mass, which resembles the gut of *X. tropicalis* at *NF 41* (cf. Figs. 2B and 5A). The anteriormost portion of the gut is located on the embryo’s left side but then bends right, forming the gastroduodenal loop (GD). Anterior to the loop, however, the foregut is much larger than in *X. tropicalis*. Posterior to the GD, the gut first straightens to form the midgut and then transitions to the hindgut caudally (Fig. 1B). At the cellular level, the gut tube comprises a mass of dense, yolky endodermal cells that surrounds a central lumen. Immediately ventral to the foregut, the liver is distinguishable by the diverticulum (Fig. 5A, B).

**Figure 5.**
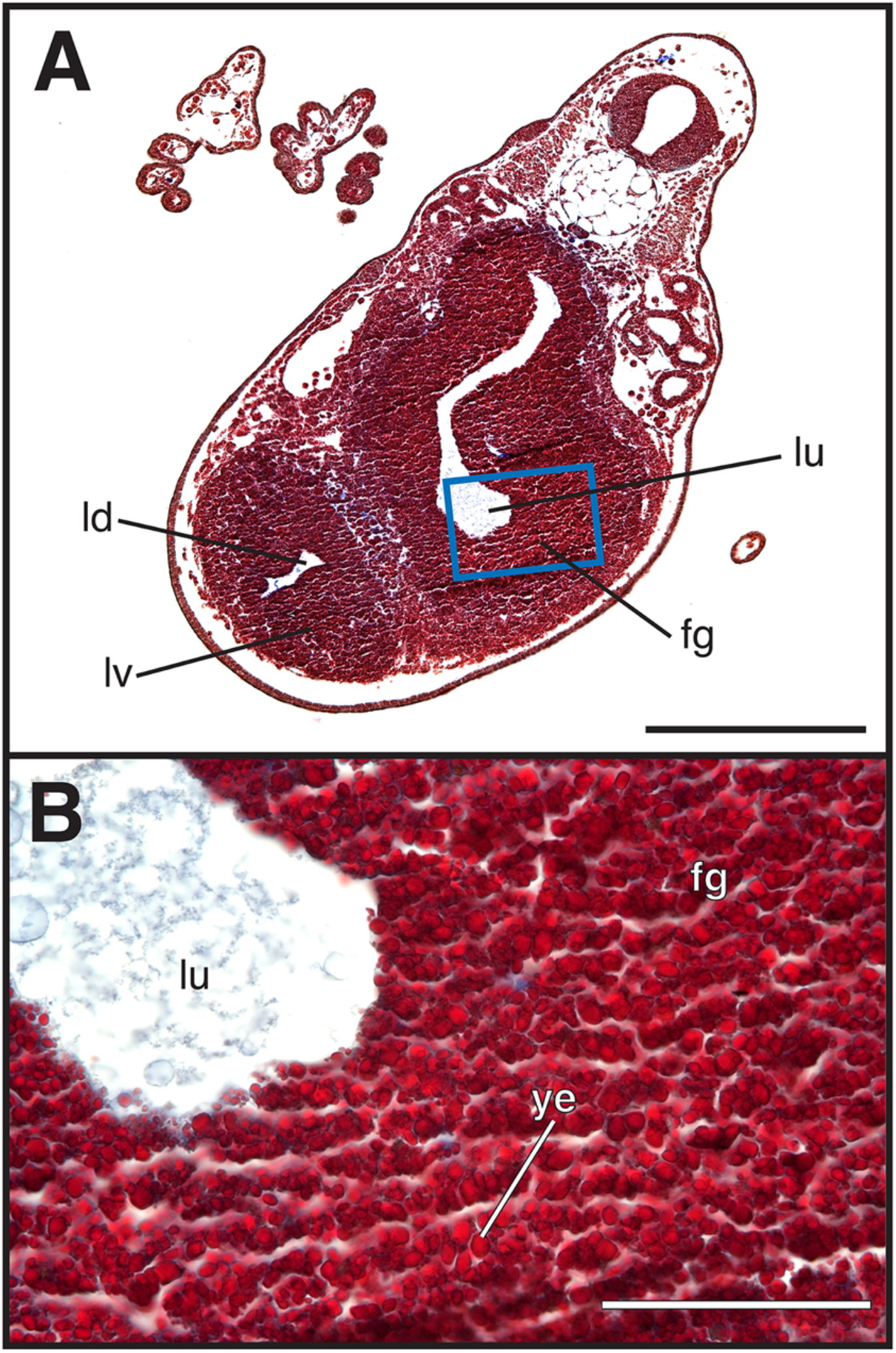
*Lepidobatrachus laevis* embryo, *GS 21*. A) Transverse section; dorsal is in upper right The foregut (fg) is recognized by its central lumen (lu); the liver (lv) is discernible by its diverticulum (ld). Area enclosed by the blue rectangle is enlarged in B. B) The foregut (fg) consists of yolky endodermal cells (ye) surrounding the lumen (lu). Magnification: A, 10X; B, 40X. Scale bar: A, 500 μm; B, 100 μm.

At *GS 23*, the gut has elongated and twists more extensively (Fig. 1B). After descending caudally from the oesophagus on the embryo’s left side, the stomach is situated ventrally and bends transversely to the right. Posterior to the stomach, the gut transitions to the intestine and bends dorsally, then descends caudally on the embryo’s right before bending transversely left. Posteriorly, the gut bends dorsally, becoming the proctodeum, which descends caudally. The stomach has begun to differentiate into distinct regions. In the anterior of the stomach is the presumptive fundic region, which transitions to the presumptive pyloric region as the stomach descends ventrally (Fig. 6A). As the stomach transitions to the intestine, the lumen ends and the intestine remains filled with yolky cells. Gastric glands are beginning to form in the anterior foregut, or presumptive fundic stomach, but the epithelium remains yolky (Fig. 6B). In the presumptive pyloric stomach, the epithelium is also quite yolky but a thin submucosa layer and a thick band of muscle have begun to differentiate (Fig. 6C). The presumptive intestine remains a solid mass of yolky, undifferentiated cells (Fig. 6D).

**Figure 6.**
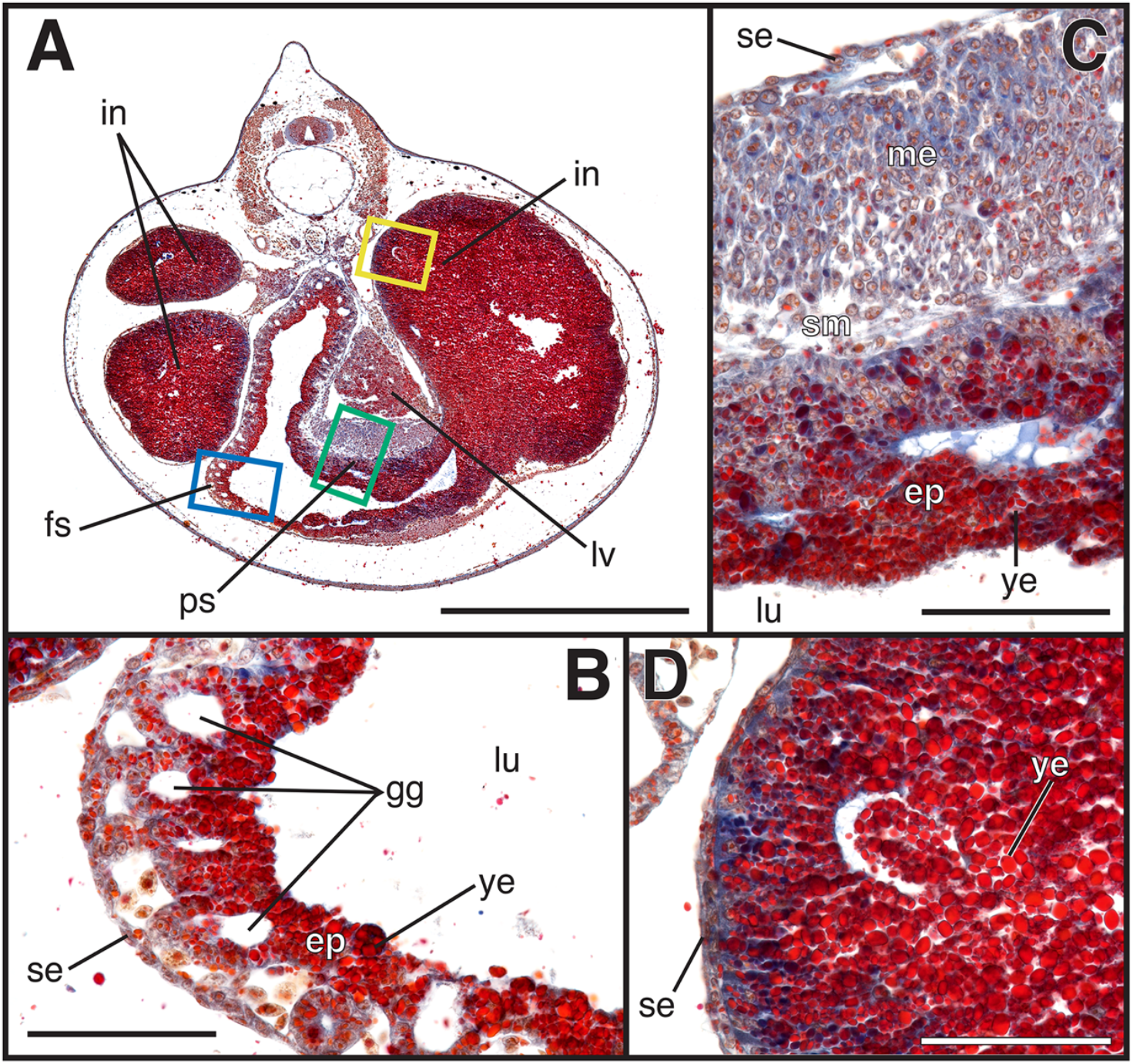
*Lepidobatrachus laevis* embryo, *GS 23*. A) Transverse section; dorsal is at the top. The fundic stomach (fs) is in the center of the visceral cavity and transitions into the pyloric stomach (ps) as the gut loops ventrally. The pyloric stomach then transitions into the intestine (in) as the gut loops to the embryo’s right side. Two more intestinal loops are visible to the left of the stomach. The liver (lv) is to the right of the stomach. Areas enclosed by the blue, green and yellow rectangles are enlarged in B, C and D, respectively. B) In the fundic stomach, an outer serosa (se) is developing while the epithelium (ep) remains mostly yolky endodermal cells (ye) surrounding the lumen (lu). Gastric glands (gg) are forming between these layers. C) The pyloric stomach has an epithelium (ep) consisting of yolky endodermal cells (ye) facing the lumen (lu). A submucosa (sm) and muscularis externa (me) are beginning to form external to the epithelium, and in turn are surrounded by an outer serosa (se). D) The intestine has an outer serosa (se), while the interior comprises a solid mass of yolky endoderm (ye). Magnification: A, 10X; B–D, 40X. Scale bar: A, 1000 μm; B–D, 100 μm.

*Lepidobatrachus laevis* larvae hatch at *GS 25* and begin feeding ravenously almost immediately. The topology of the gut is similar to *GS 23* (Fig. 1B). The stomach has shifted to the larva’s right side and is much thicker and more voluminous. The intestine has elongated, although much less than in *X. tropicalis*, and loops ventrally next to the stomach on the left side. The entire gastrointestinal tract is well differentiated regionally, with the stomach occupying most of the visceral cavity centrally. Loops of the intestine are situated dorsal to the stomach on the right and dorsal and lateral to it on the left (Fig. 7A). Beginning in the oesophagus, the epithelium consists of a single layer of ciliated columnar cells. The submucosa consists of loose connective tissue and is surrounded by a layer of muscularis externa so thin it is barely distinguishable from the serosa (Fig. 7B). The fundic stomach is the largest portion of the stomach by volume. The epithelium is simple cuboidal,; it is interrupted periodically by goblet cells and gastric pits. The mucosa is thin with little connective tissue but is full of tubular gastric glands (Fig. 7C). In the pyloric stomach, the epithelium is a folded layer of simple columnar cells. There is no muscularis mucosa, but basal to the epithelium is a thick submucosa. It, in turn, is surrounded by the thick muscularis externa (Fig. 7D). Finally, the serosa is the outermost layer of the stomach. In contrast, the intestinal lining is still very yolky and does not yet have a distinct epithelium (Fig. 7A).

**Figure 7.**
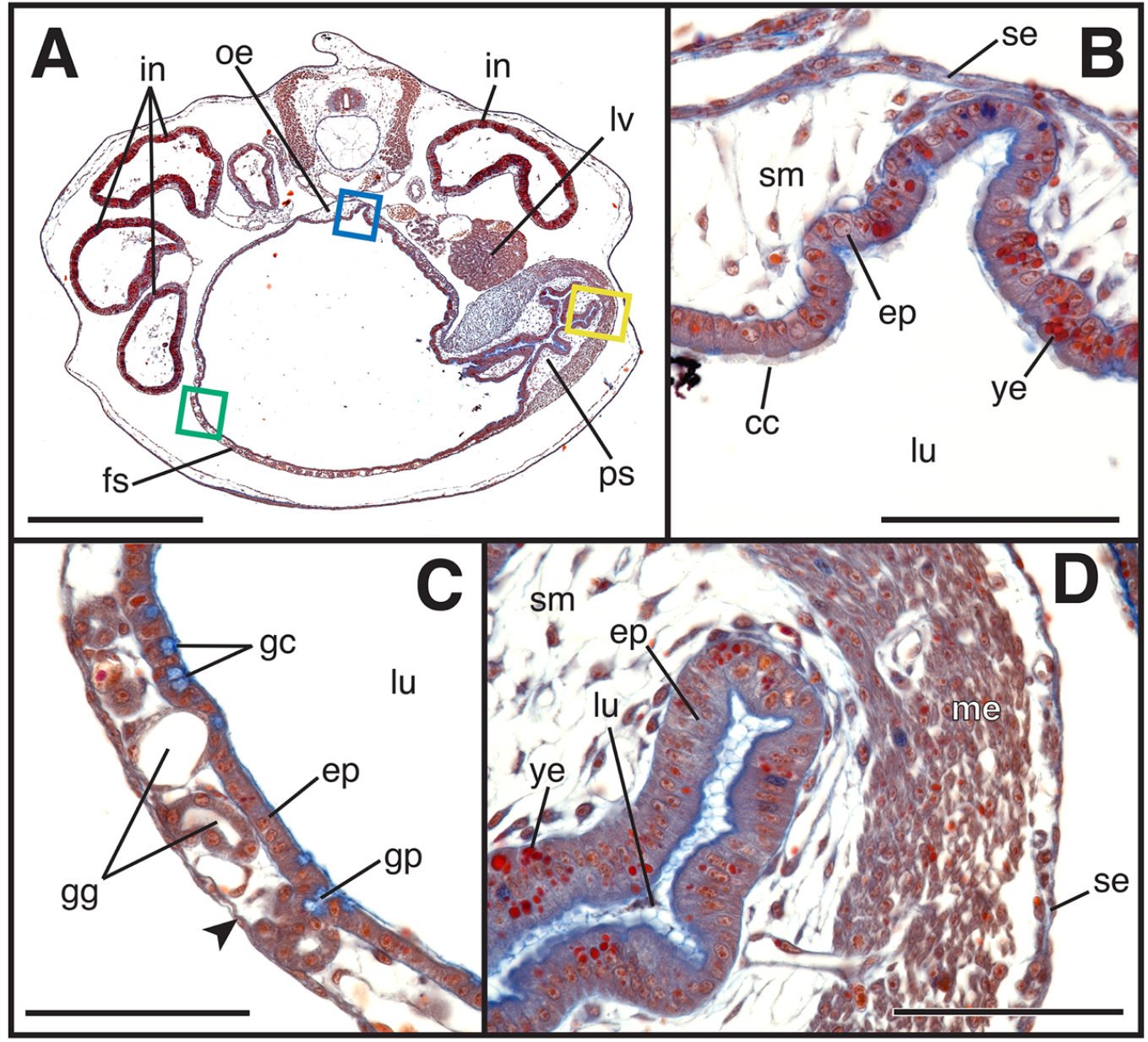
*Lepidobatrachus laevis* tadpole, *GS 25*. A) Transverse section; dorsal is at the top. The fundic stomach (fs) dominates the visceral cavity and is continuous with the oesophagus (oe) dorsally and the pyloric stomach (ps) ventrally and to the right. Areas enclosed by the blue, green and yellow rectangles are enlarged in B, C and D, respectively. The liver (lv) is to the right and the intestines (in) loop to either side of the stomach. B) The oesophagus has a simple cuboidal epithelium (ep) with ciliated cells (cc) facing the lumen (lu) but retains a few patches of yolky endoderm (ye). External to the epithelium is the submucosa (sm), which in turn is invested by the serosa (se). C) The fundic stomach consists of a simple cuboidal epithelium (ep) facing the lumen (lu), a middle layer containing gastric glands (gg), and an outermost layer comprising a very thin muscularis externa and adjacent serosa (arrowhead). The epithelium contains mucous goblet cells (gc) and gastric pits (gp). D) The pyloric stomach consists of a simple columnar epithelium (ep) facing the lumen (lu), submucosa (sm), muscularis externa (me), and outermost serosa (se). A few yolky endodermal cells (ye) remain in the epithelium. Magnification: A, 10X; B–D, 40X. Scale bar: A, 1000 μm; B–D, 100 μm.

### Mid-larval stomach development

#### Xenopus tropicalis

At *NF 53*, the topology of the gut is little changed from *NF 45*, although the intestine has elongated and formed additional coils. The stomach is slightly thicker than the intestine, both in overall diameter and in thickness of the gut wall, but it is very short in comparison to the length of the intestine (Fig. 8A). The same essential components of the stomach present at *NF 45* are also observed at *NF 53*, but the microanatomy of the stomach is now more complex. The epithelium is more distinct and consists of pseudostratified cells capped with blue-staining mucus and folded into gastric pits. The mucosa contains more elaborate tubular gastric glands (Fig. 8B). There is still no muscularis mucosa or submucosa. The intestinal epithelium is also coated in mucus apically, with a thin layer of submucosa enclosed by the serosa (Fig. 8C).

**Figure 8.**
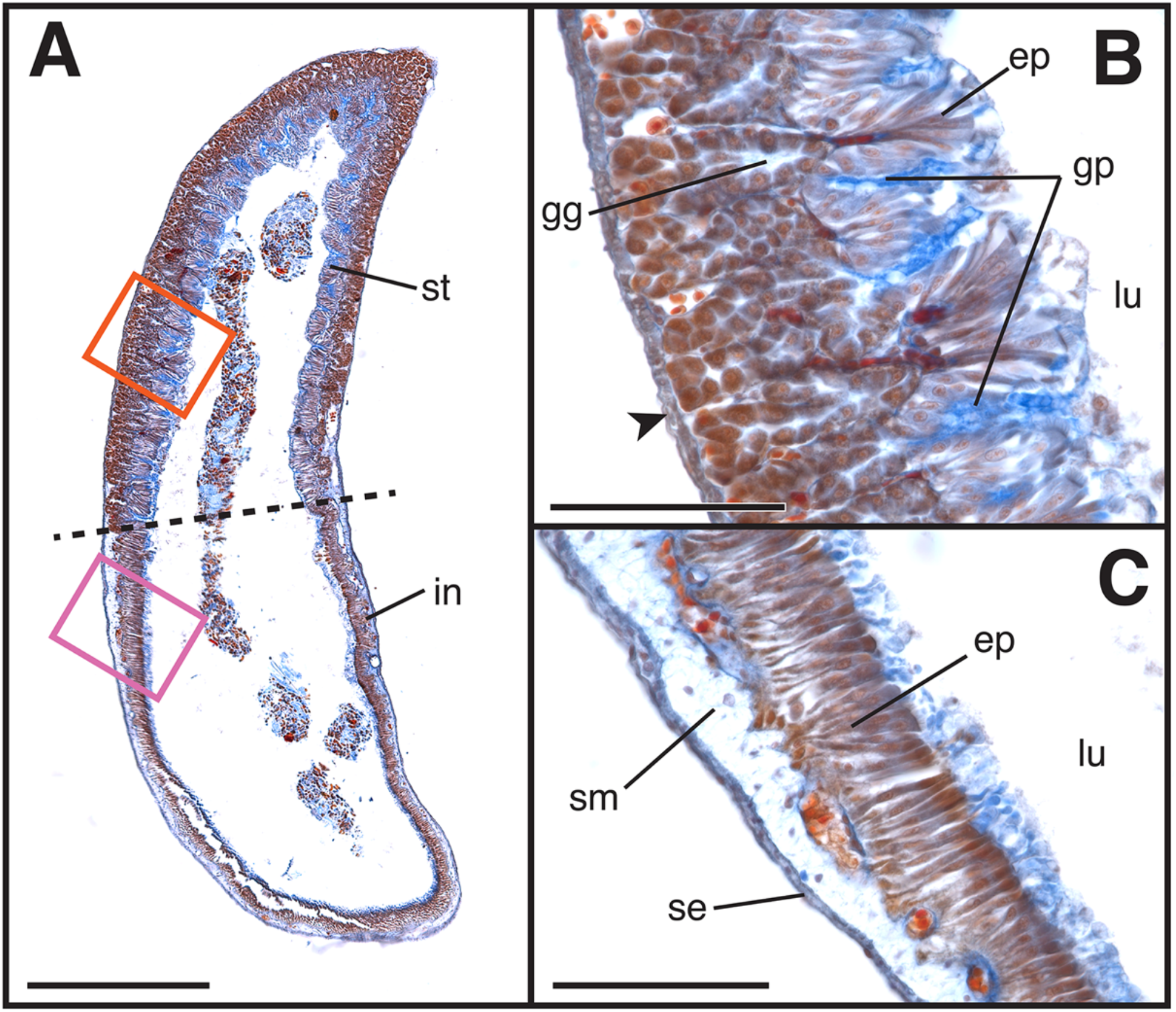
Stomach of *Xenopus tropicalis* tadpole, *NF 53*. A) Oblique section through the gut tube in the region where the stomach (st) transitions into the intestine (in). Dashed line indicates the transition zone. Areas enclosed by the orange and pink rectangles are enlarged in B and C, respectively. B) The stomach wall comprises an inner epithelium (ep) and an outer layer of muscularis externa and serosa (arrowhead); the two regions are separated by a thick layer of gastric glands (gg). The epithelium faces the lumen (lu) and is interspersed with gastric pits (gp). C) In the intestine, the inner epithelium (ep) facing the lumen (lu) is separated from the outer serosa (se) by the submucosa (sm), which lacks gastric glands. Magnification: A, 10X; B–C, 40X magnification. Scale bar: A, 500 μm; B–C, 100 μm.

#### Lepidobatrachus laevis

The topology of the gut at *GS 31* is very similar to *GS 25*, but the stomach is disproportionately much larger. The intestine, while slightly longer, is not coiled nearly as extensively as it is in *X. tropicalis*. The microanatomy of the stomach, while similar to *GS 25*, is much more elaborate. The fundic stomach is anteriormost, followed by the pyloric stomach and the intestine (Fig. 9A). The fundic epithelium has a brush border and is dotted with many goblet cells (Fig. 9B). The mucosa has many more gastric glands containing mucous neck cells embedded in the lamina propria. In the pyloric region, the epithelium folds into gastric pits (Fig. 9C). The deep connective tissue has a dense layer adjacent to the epithelium and a looser layer adjacent to the muscularis externa, possibly representing the initial differentiation of distinct lamina propria and submucosa, although still lacking a muscularis mucosa. The muscularis externa is also thicker and denser, with an inner circular layer and an outer longitudinal layer.

**Figure 9.**
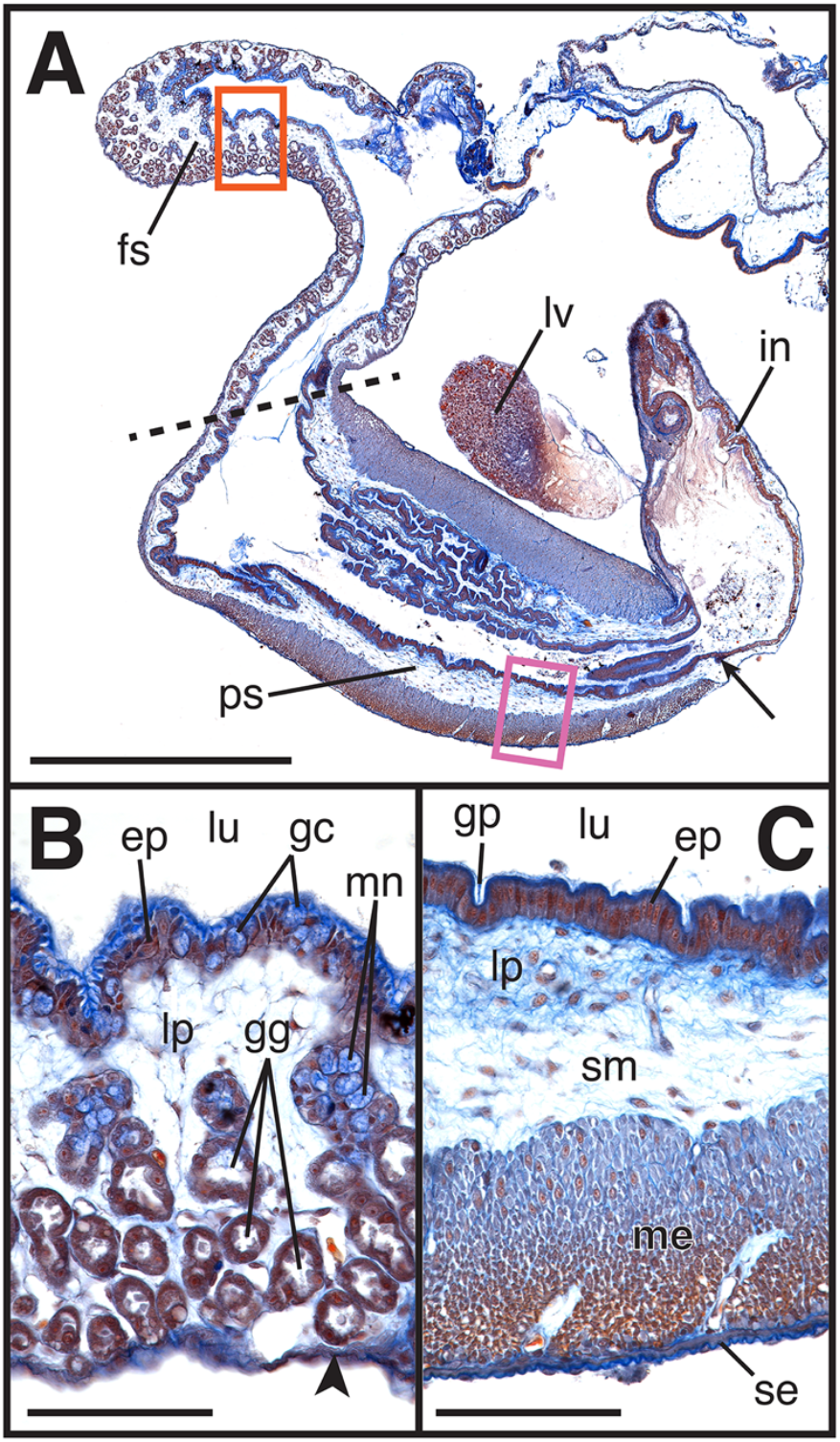
Stomach of *Lepidobatrachus laevis* tadpole, *GS 31*. A) Longitudinal section through the stomach; anterior is at the top. The larval stomach consists of a fundic portion (fs) anteriorly and a pyloric portion (ps) posteriorly. Dashed line indicates the transition zone. The pyloric stomach narrows posteriorly into a sphincter (arrow) that opens into the intestine (in). The liver (lv) lies immediately dorsal to the pyloric stomach. Areas enclosed by the orange and pink rectangles are enlarged in B and C, respectively. B) The epithelium (ep) of the fundic stomach is densely populated with goblet cells (gc), which face the lumen (lu). The adjacent lamina propria (lp) is thick and full of gastric glands (gg), which contain mucous neck cells (mn). The thin muscularis externa and serosa (arrowhead) are the outermost layers. C) The epithelium (ep) of the pyloric stomach faces the lumen (lu) and contains gastric pits (gp). The adjacent lamina propria (lp) is successively enclosed by the submucosa (sm), then a thick muscularis externa (me) and finally the outermost serosa (se). Magnification: A, 10X; B–C, 40X. Scale bar: A, 1000 μm; B–C, 100 μm.

### Stomach Development During Metamorphosis

#### Xenopus tropicalis

At *NF 60*, metamorphic climax has begun (Ishizuya-Oka & Ueda, 1996; Ishizuya-Oka *et al*., 1998; Schrieber *et al*., 2004). At a gross level, the configuration of the digestive tract is still similar to *NF 53*, but the microanatomy has begun to change (Fig. 10A). The epithelium of the larval stomach is the most apical layer of the stomach, it stains less vibrantly, and cell morphology is less distinct. While this “larval epithelium” remains attached to the basement membrane in some areas (Fig. 10B, C), it has begun to detach in others (Fig. 10D). The basement membrane has thickened and stains dark blue, separating the larval epithelium from adult epithelium. Basal to the larval epithelium is developing a second, “adult epithelium,” indicated by the presence of a muscularis externa, a layer of muscle between the glandular mucosa and the serosa, and new gastric glands (Fig. 10B-D).

**Figure 10.**
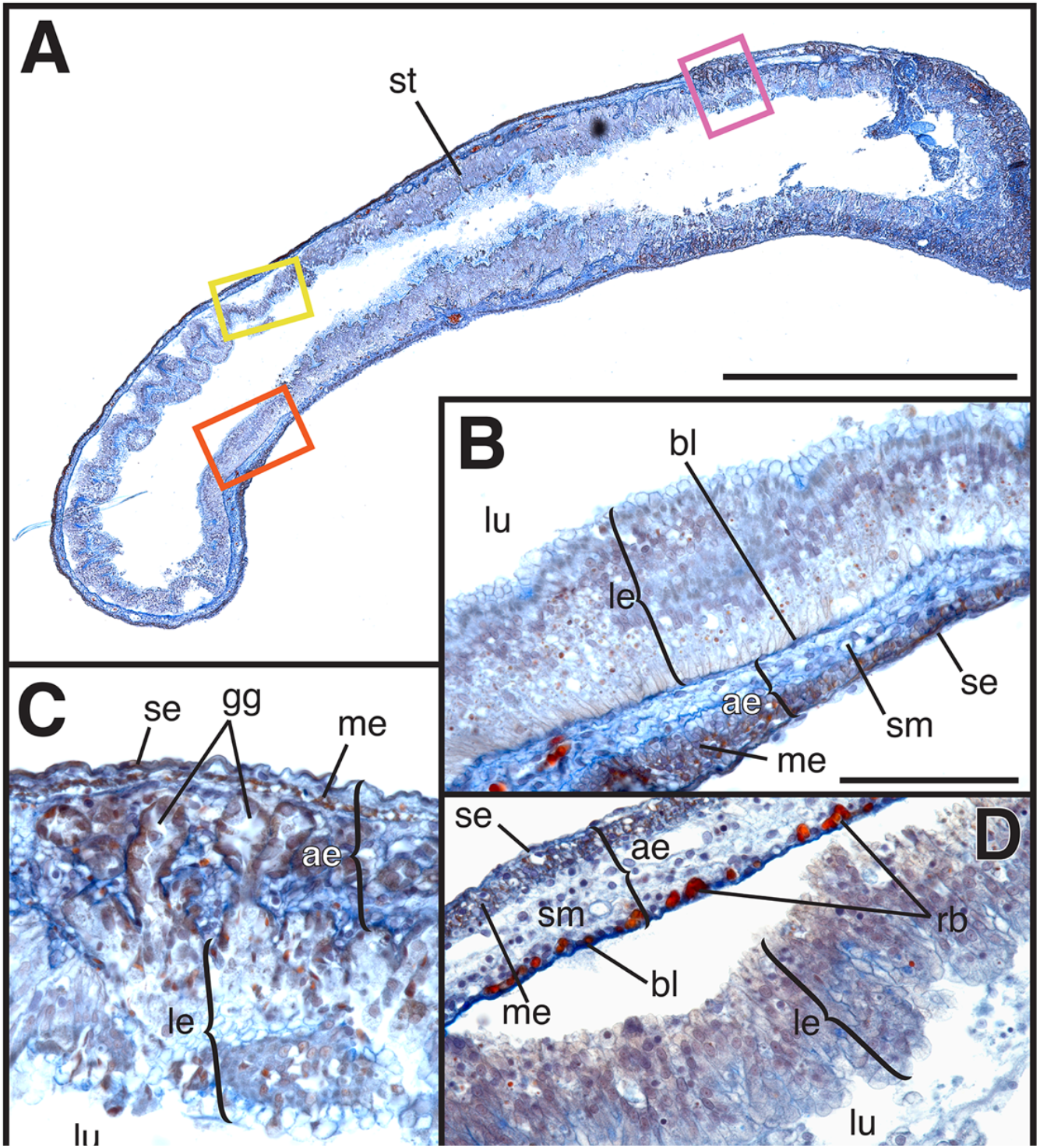
Stomach of metamorphosing *Xenopus tropicalis, NF 60*. A) Oblique section through the stomach (st), positioned with anterior to the right and posterior to the left. Areas enclosed by the orange, pink and yellow rectangles are enlarged in B, C and D, respectively. B) The adult epithelium (ae) has formed basal to the larval epithelium (le); the two layers are separated by a thick basal lamina (bl). Exterior to the basal lamina is the submucosa (sm) followed by a muscularis externa (me) and an outermost serosa (se). C) The larval epithelium (le) faces the lumen (lu) and overlies the adult epithelium (ae), which is developing gastric glands (gg) and is surrounded externally by the developing muscularis externa (me) and serosa (se). D) The larval epithelium (le) has detached from the adult epithelium (ae) and entered the lumen (lu). Deep to the detaching larval epithelium, the innermost layer of the adult stomach consists of a basal lamina (bl) followed by red blood cells (rb), submucosa (sm), muscularis externa (me) and outermost serosa (se). Magnification: A, 10X; B–D, 40X. Scale bar: A, 1000 μm; B–D, 100 μm.

By *NF 63*, the stomach has begun to expand and the intestine is shortening. Throughout the gastrointestinal tract, the larval epithelium is no longer present; only the developing adult epithelium remains. The stomach has differentiated into two distinct regions with contrasting morphology: a fundic region anteriorly and a pyloric region posteriorly (Fig. 11A). In the fundic region, the epithelium is still disorganized, but a thick layer of tubular gastric glands is present external to the epithelium (Fig. 11B). The submucosa has also thickened and contains pockets of red blood cells. In the pyloric region, the epithelium comprises a single layer of columnar cells that folds into a series of gastric pits (Fig. 11C). Basal to this, the submucosa is dense and the muscularis externa is thicker than in the fundic region. The serosa remains the outermost layer of both regions.

**Figure 11.**
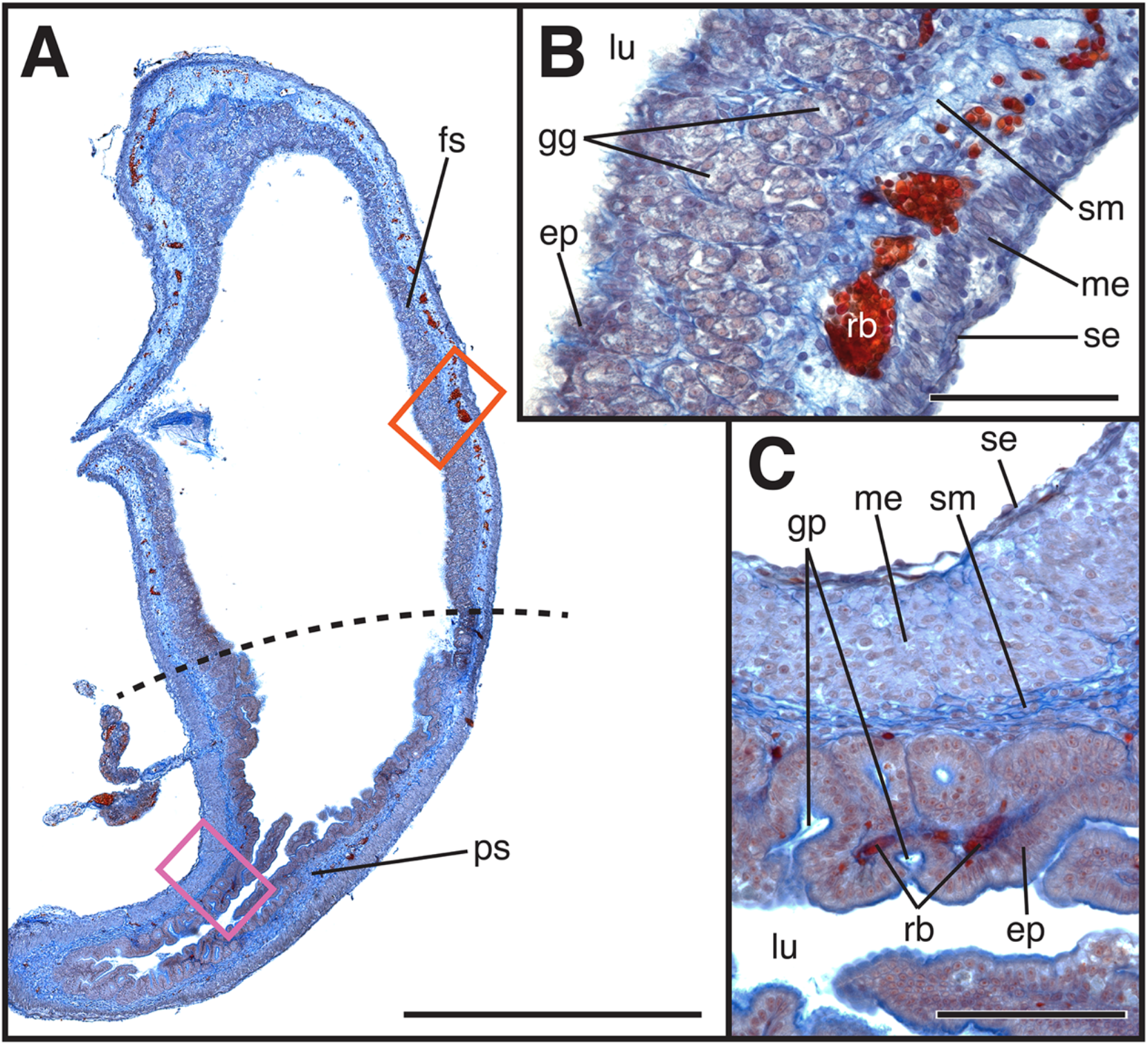
Stomach of metamorphosing *Xenopus tropicalis, NF 63*. A) Oblique section through the stomach showing the fundic stomach (fs) and pyloric stomach (ps), which are demarcated by a dashed line. Areas enclosed by the orange and pink rectangles are enlarged in B and C, respectively. B) Internally, the fundic stomach consists of an epithelium (ep) facing the lumen (lu). External to this is a layer of gastric glands (gg) followed by the submucosa (sm), which contains patches of red blood cells (rb). Surrounding this are the muscularis externa (me) and an outermost serosa (se). C) In the pyloric stomach, the epithelium (ep) facing the lumen (lu) is folded into gastric pits (gp), with pockets of red blood cells (rb) in the submucosa (sm) just external to the epithelium. Enclosing these layers are the muscularis externa (me) and an outermost serosa (se). Magnification: A, 10X; B–C, 40X;. Scale bar: A, 1000 μm; B–C, 100 μm; C.

At NF 65, immediately before the end of metamorphosis, the stomach is considerably larger and the intestine is much shorter than in earlier larval stages; the intestine is reduced to just one or two coils. The stomach has even more apparent regionalization of fundic and pyloric regions (Fig. 12A). The fundic region is well differentiated, with a distinct epithelium dotted with gastric pits and subtended by gastric glands (Fig. 12B). The submucosa surrounds the glandular mucosa, which in turn is surrounded by the muscularis externa and outermost serosa. In the pyloric region, the epithelium faces the lumen and is surrounded by dense connective tissue of the lamina propria, which in turn is enclosed by the looser connective tissue of the submucosa (Fig. 12C). There still appears to be no muscularis mucosa. External to this is the muscularis externa and, finally, the serosa.

**Figure 12.**
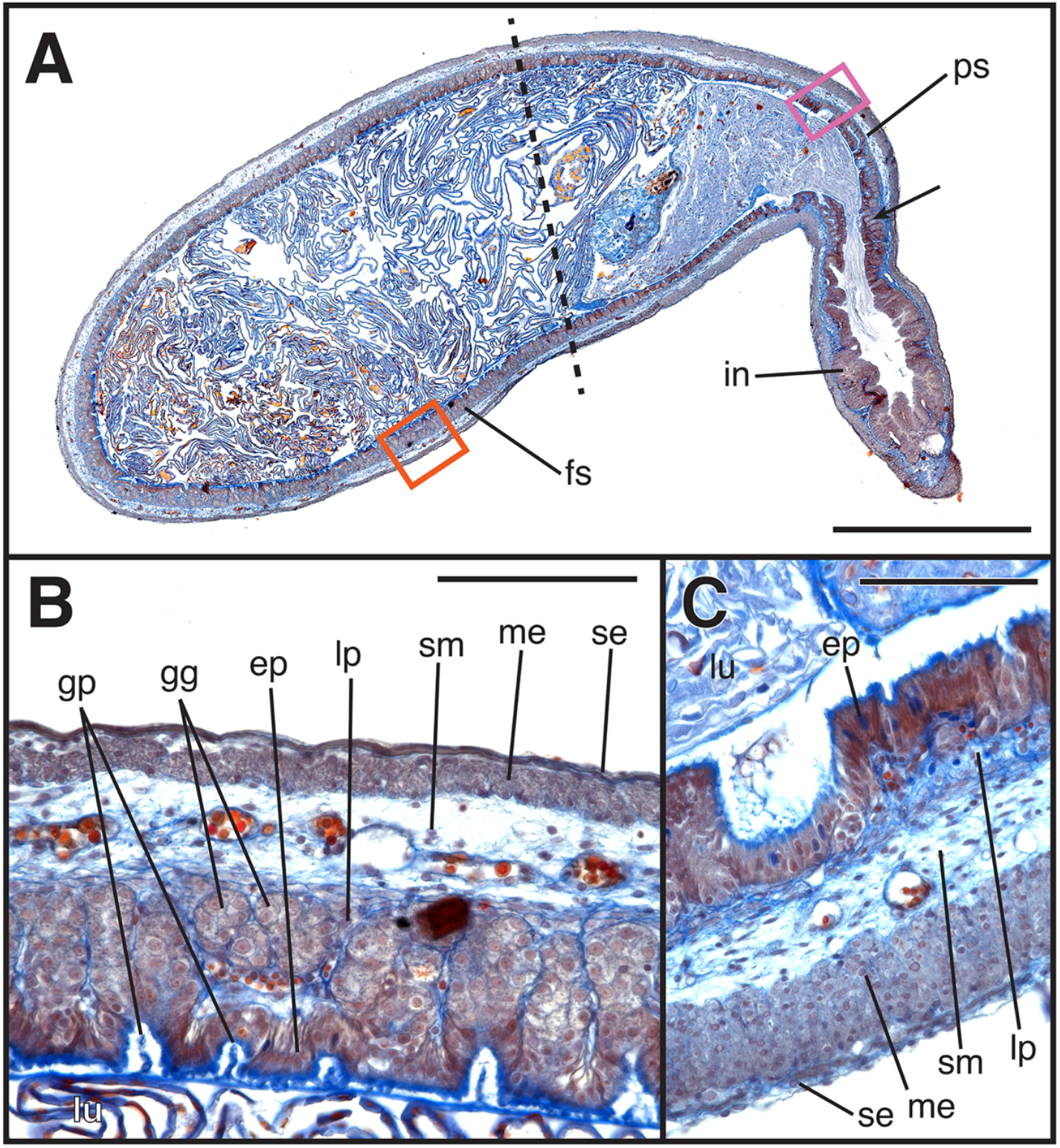
Stomach of adult *Xenopus tropicalis, NF 65*. A) The stomach comprises two distinct morphological regions: the fundic stomach (fs) anteriorly, followed by the pyloric stomach (ps) posteriorly; the two regions are demarcated by a dashed line. The intestine (in) is separated from the pyloric stomach by the pyloric sphincter (arrow). Areas enclosed by the orange and pink rectangles are enlarged in B and C, respectively. B) In the fundic stomach, the epithelium (ep) faces the lumen (lu) and has numerous gastric pits (gp) and the lamina propria (lp) contains gastric glands (gg). External to the glands are the submucosa (sm), muscularis externa (me), and the outermost serosa (se). C) In the pyloric stomach, the epithelium (ep) is the innermost layer facing the lumen (lu), followed by the lamina propria (lp), the submucosa (sm), a thick muscularis externa (me), and finally the serosa (se). Magnification: A, 10X; B–C, 40X. Scale bar: A, 1000 μm; B–C, 100 μm.

#### Lepidobatrachus laevis

The topology of the gut does not change during metamorphosis, when froglets continue to feed as aggressively as do tadpoles, if not more so. Stomach microanatomy at *GS 41* looks very similar to that at *GS 31*, with only a few minor differences. The connective tissue is denser throughout the stomach (Fig. 13A). In the transitional region between the oesophagus and the fundic stomach (sometimes called the cardiac stomach), the gastric glands contain many mucous neck cells (Fig. 14B). In the posterior fundic region, tubular gastric glands are even more abundant and the epithelium is more highly folded with gastric pits (Fig. 14C). Throughout the fundic region, the muscularis externa has thickened and is now distinct from the serosa (Fig. 13B-C). In the pyloric region, the gastric pits in the epithelium have deepened. The lamina propria has now fully differentiated and is separated from the submucosa by the muscularis mucosa (Fig. 13D).

**Figure 13.**
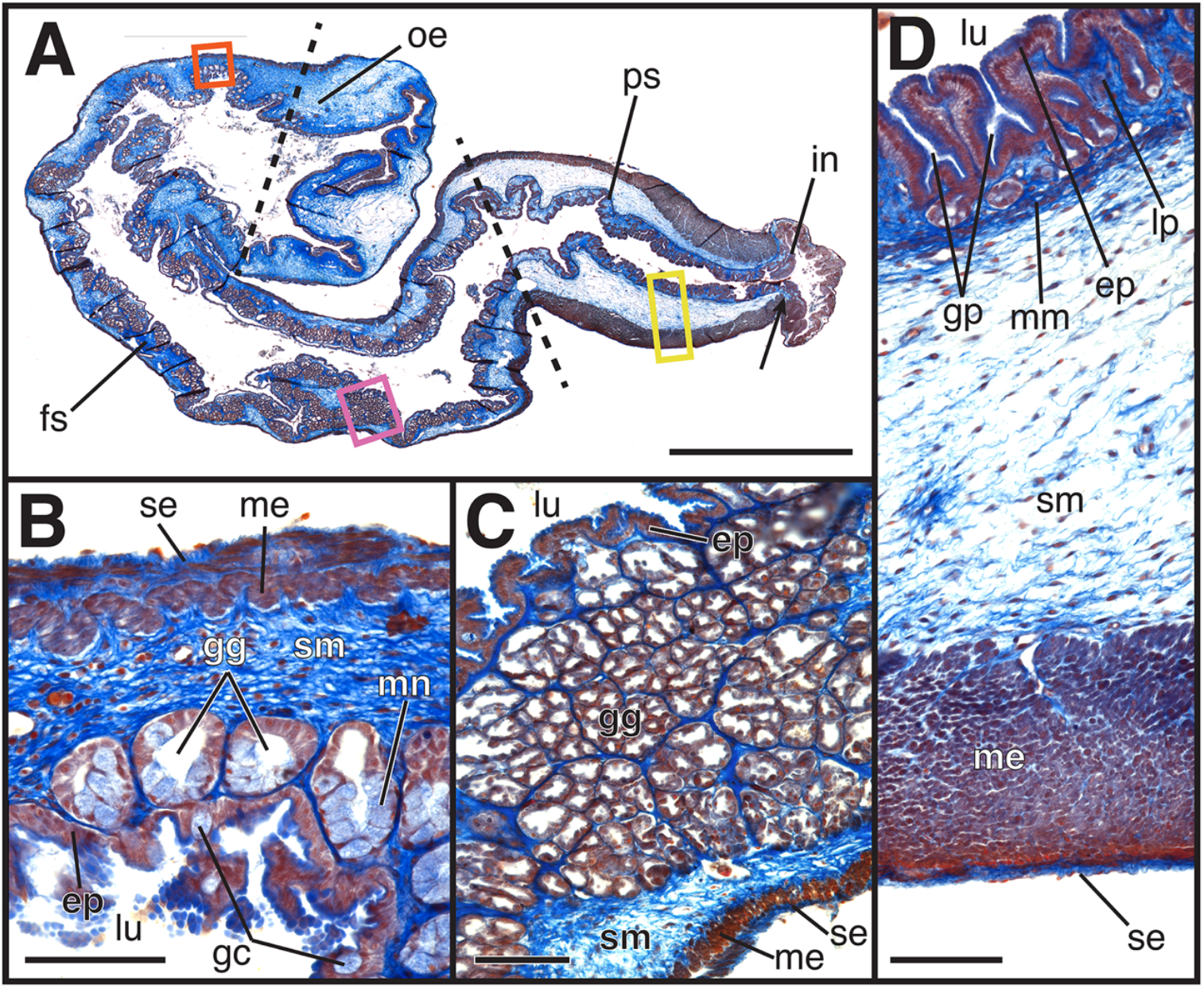
Stomach of metamorphosing *Lepidobatrachus laevis, GS 41*. A) Longitudinal section of the stomach showing the oesophagus (oe), fundic stomach (fs) and pyloric stomach (ps), which are demarcated by dashed lines. The intestine (in) is separated from the pyloric stomach by the pyloric sphincter (arrow). Areas enclosed by the orange, pink and yellow rectangles are enlarged in B, C and D, respectively. B) In the anterior portion of the fundic stomach, the epithelium (ep) faces the lumen (lu) and is highly folded and spotted with goblet cells (gc). Deep to this layer are gastric glands (gg) filled with mucous neck cells (mc), followed by the submucosa (sm), a thin muscularis externa (me) and the serosa (se). C) In the posterior fundic stomach, the epithelium (ep) faces the lumen (lu) and also is highly folded and even more densely populated with gastric glands (gg). Deep to the glands are the submucosa (sm), muscularis externa (me) and serosa (se). D) The pyloric stomach has a highly folded epithelium (ep) facing the lumen (lu) with numerous, deep gastric pits (gp). Basal to the epithelium is the lamina propria (lp), followed by a thin muscularis mucosa (mm), a thick submucosa (sm), a thick muscularis externa (me) and finally the serosa (se). Magnification: A, 5X; B–D, 40X. Scale bar: A, 2000 μm; B–D, 100 μm.

**Figure 14.**
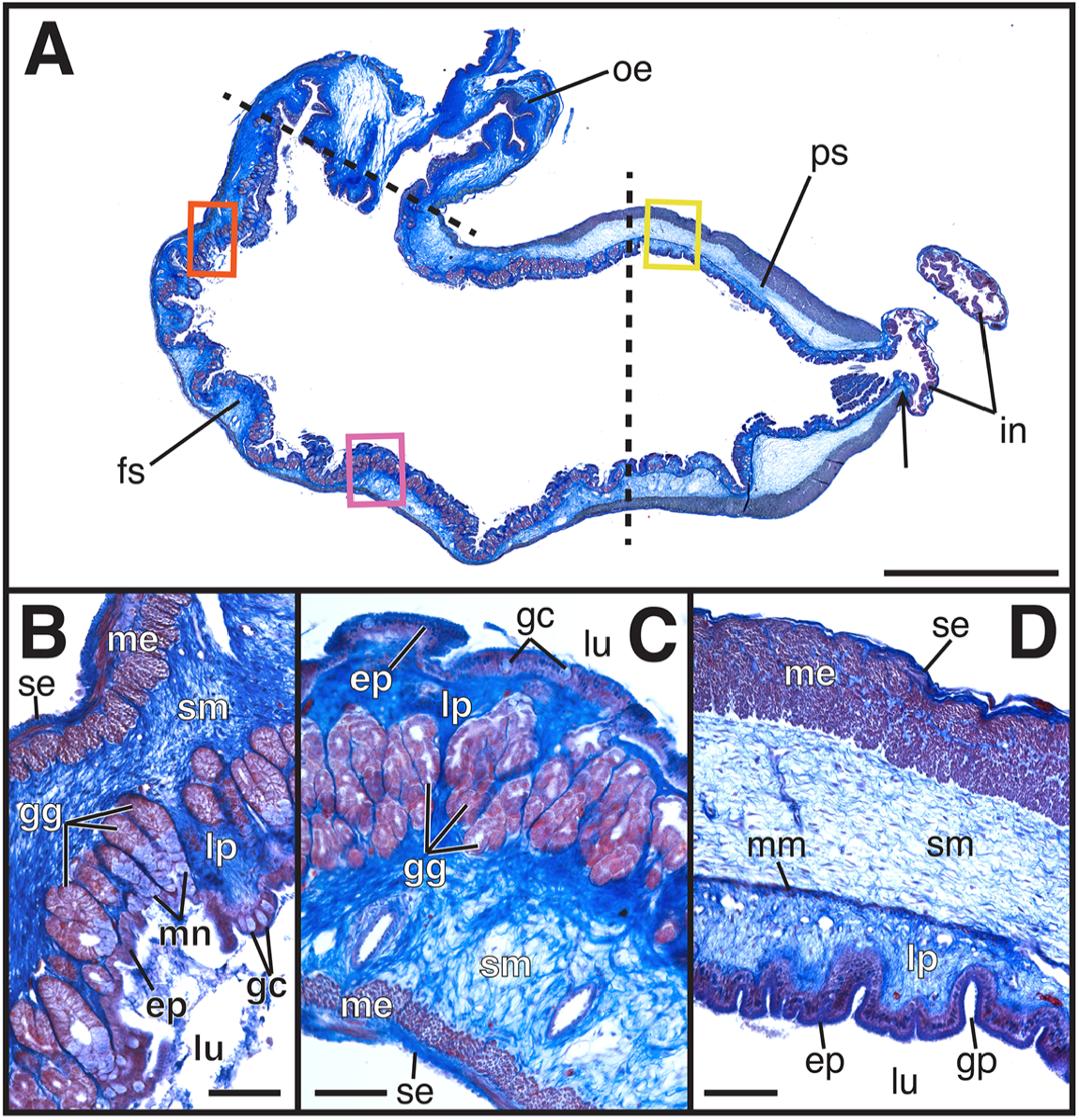
Stomach of metamorphosing *Lepidobatrachus laevis, GS 44*. A) Longitudinal section showing the oesophagus (oe), fundic stomach (fs) and pyloric stomach (ps), which are demarcated by dashed lines. The intestine (in) is separated from the pyloric stomach by the pyloric sphincter (arrow). Areas enclosed by the orange, pink and yellow rectangles are enlarged in B, C and D, respectively. B) In the anterior portion of the fundic stomach, the epithelium (ep) is the most apical layer and occasionally contains goblet cells (gc). Basal to this layer is the lamina propria (lp), which contains gastric glands (gg), many filled with mucous neck cells (mc) that empty into the lumen (lu). The lamina propria is enclosed by the submucosa (sm), the muscularis externa (me) and finally the serosa (se). C) In the posterior fundic stomach, the epithelium (ep) faces the lumen (lu). The epithelium also contains sporadic goblet cells (gc) and is highly folded. The lamina propria (lp) is densely populated with gastric glands (gg). Basal to the glands are the submucosa (sm), muscularis externa (me) and serosa (se). D) The pyloric stomach has a highly folded epithelium (ep) with numerous gastric pits (gp). Deep to the epithelium is the lamina propria (lp), followed by a thin muscularis mucosa (mm), a thick submucosa (sm), a thick muscularis externa (me) and finally the serosa (se). Magnification: A, 5X; B–D, 20X. Scale bar: A, 2000 μm; B–D, 100 μm.

At *GS 44*, the stomach looks very similar to *GS 41* (Fig. 14A). Both the anterior and posterior parts of the fundic stomach appear unchanged (Fig. 14B-C). Likewise, the pyloric region is unchanged except for the continued differentiation of the muscularis mucosa (Fig. 14D).

At *GS 46*, the froglet has completed metamorphosis and continues to feed aggressively. The overall morphology of the stomach is practically identical to *GS 44* (Fig. 15A). Although the anterior portion of the fundic stomach still lacks a muscularis mucosa (Fig. 15B), it can now be seen separating the glandular lamina propria from the aglandular submucosa in the posterior region of the fundic stomach (Fig. 15C). The pyloric stomach is unchanged (Fig. 16D).

**Figure 15.**
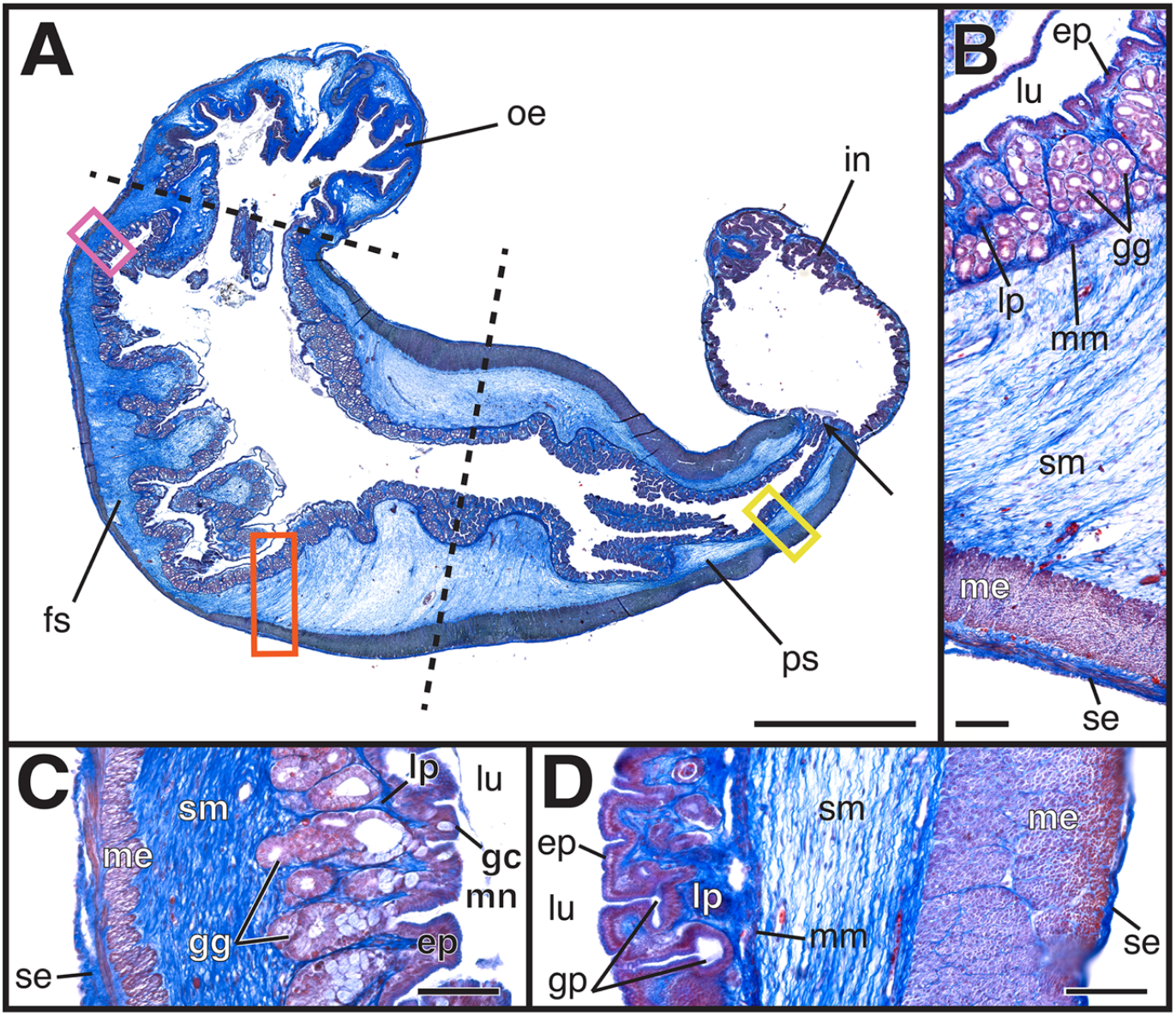
Stomach of an adult *Lepidobatrachus laevis, GS 46*. A) Longitudinal section showing the oesophagus (oe), fundic stomach (fs) and pyloric stomach (ps), which are demarcated by dashed lines. The intestine (in) is separated from the pyloric stomach by the pyloric sphincter (arrow). Areas enclosed by the orange, pink and yellow rectangles are enlarged in B, C and D, respectively. B) In the posterior fundic stomach, the epithelium (ep) faces the lumen (lu). Basal to the epithelium, the lamina propria (lp) is densely populated with gastric glands (gg) and is separated from the submucosa (sm) by the thin muscularis mucosa (mm). Basal to the submucosa (sm) are the muscularis externa (me) and serosa (se). C) In the anterior portion of the fundic stomach, the epithelium (ep) is the most apical layer and occasionally contains goblet cells (gc). Basal to this layer is the lamina propria (lp), containing gastric glands (gg), many filled with mucous neck cells (mc) that empty into the lumen (lu). The mucosa of the gastric glands is followed by the submucosa (sm), the muscularis externa (me) and the serosa (se). D) The pyloric stomach has a highly folded epithelium (ep) with numerous gastric pits (gp). Deep to the epithelium is the lamina propria (lp), followed by a thin muscularis mucosa (mm), a thick submucosa (sm), a thick muscularis externa (me) and finally the serosa (se). Magnification: A, 5X; B–D, 20X. Scale bar: A, 2000 μm; B–D, 100 μm.

## Discussion

### *The larval stomach of* Lepidobatrachus laevis *is adult-like*

The larval stomach of *Lepidobatrachus laevis* is highly unusual among metamorphosing frogs. The only other tadpole known to have a similar stomach and developmental trajectory belongs to the congeneric species *L. llanensis*, and to a lesser degree frogs of the genus *Ceratophrys* (Fabrezi & Cruz, 2020). (Larval stomach morphology of the third species of *Lepidobatrachus, L. asper*, is not described, but presumably it resembles that of its two congeners.) The stomach of typical tadpoles is adapted to a microphagous, herbivorous diet: it’s very small, lacks anatomical subfunctionalization, and does not secrete hydrochloric acid or pepsinogen, two digestive compounds found in stomachs of adult frogs (Barrington, 1945; Lambertini, 1946; Dodd & Dodd, 1976; Fox, 1983; Griffiths, 1961). The stomach of larval *Lepidobatrachus* instead is suited to digest a megalophagous, carnivorous diet: it’s very large, comprises distinct glandular fundic and muscular pyloric regions, and secretes both hydrochloric acid and pepsinogen (Carrol *et al*., 1991). It is in all significant respects an adult-like stomach in a larva.

Here we confirm the developmental pattern of the larval stomach reported in earlier studies of *L. laevis* (Ruibal & Thomas, 1988; Bloom, 2013; Amin, 2015; Fabrezi & Cruz, 2020), but we also expand on these works by describing embryonic development of the stomach at Gosner stages 21, 23 and 25, as well as throughout metamorphosis at GS 41, 44 and 46. While the gastrointestinal tract develops initially as an undifferentiated yolky tube, the foregut soon forms a distinct anterior, glandular region and a posterior, muscular region, both of which are present before hatching. This ontogenetic sequence differs significantly from the stereotypical, biphasic pattern of stomach development seen in most metamorphosing anurans such as *X. tropicalis*, in which the larval stomach is very small and contains a homogeneous glandular mucosa but lacks a muscular region. Additionally, the larval gastric epithelium in *L. laevis* is functionally more complex in containing mucin-producing goblet cells and mucous neck cells, which are absent from the larval epithelium in *X. tropicalis*. In sum, the adult-like stomach of larval *L. laevis* develops *de novo* during embryogenesis and is not preceded by the specialized larval stomach seen in most other metamorphosing frogs. Larval-specific features are not expressed during embryogenesis, and the developing stomach assumes a largely adult-like configuration from the outset. Consequently, even before the onset of feeding, the larval stomach of *L. laevis* more closely resembles the stomach characteristic of adult frogs than it does the larval stomach of most other metamorphosing anurans. It does, however, bear some resemblance to the larval stomach of other carnivorous tadpoles, such as the closely related *Ceratophrys*, which independently evolved larval carnivory, although in a less extreme form designated “macrophagous feeding” (Fabrezi *et al*., 2016; Fabrezi & Cruz, 2020). *Ceratophrys* also have discrete fundic and pyloric regions of the stomach as tadpoles; however, their stomachs are smaller at the onset of larval feeding and have less-folded epithelia, smaller layers of connective tissue and very little muscle (Bloom *et al*., 2013; Fabrezi & Cruz, 2020).

Despite the adult-like feeding behavior of larval *Lepidobatrachus* and the adult-like subfunctionalization of its gastric anatomy, its stomach lacks some of the complexity typical of the adult stomach of anurans and other vertebrates (Carroll, *et al*. 1991; Fabrezi & Cruz, 2020). The entire larval stomach wall, for example, lacks a middle layer of muscle, the muscularis mucosa. In the adult stomach, the muscularis mucosa subdivides adjacent connective tissue into an inner, dense, glandular lamina propria and an outer, less dense, aglandular submucosa. In the absence of the muscularis mucosa, we simply refer to connective tissue with a glandular mucosa as “lamina propria” and connective tissue without glands in the larval stomach as “submucosa.” The fundic stomach also lacks mucous neck cells in the tubular glands as well as a muscularis externa. Nevertheless, lack of these structures does not appear to inhibit larval feeding. At hatching, an *L. laevis* tadpole is capable of consuming and digesting animal prey almost as large as itself. Studies of stomach enzymatic activity also confirm that the gastric glands are capable of secreting pepsin and hydrochloric acid at the time of hatching, thus helping to digest prey (Carroll, *et al*. 1991).

During evolution of the unique larval trophic morphology of *Lepidobatrachus*, major components of adult stomach development have been shifted from metamorphosis to embryogenesis. This suggests that developmental modules that produce the typical herbivorous larval stomach phenotype have been lost from or are otherwise repressed during the ontogeny of *L. laevis*. This evolutionarily derived pattern of stomach development and its unique larval morphology underlie the ability of *L*. *laevis* to consume a similar carnivorous diet between larval and adult phases.

### *The stomach of* Lepidobatrachus laevis *does not remodel at metamorphosis*

In *L. laevis*, formation of a functional adult-like stomach by the beginning of the larval period has additional implications for the subsequent metamorphic transition from larva to adult. In metamorphosing anurans, the gastrointestinal tract typically undergoes extreme remodeling at metamorphosis (Andrew & Hickman, 1974; Rovira *et al*., 1995; Shi & Ishizuya-Oka, 1996; Schreiber *et al*., 2009). The larval epithelium degenerates, is shed into the lumen, and is replaced by an underlying adult epithelium that develops distinct fundic and pyloric regions (Ishizuya-Oka *et al*., 1998). Our observations demonstrate that, in *X. tropicalis*, stomach development during metamorphosis follows this ontogenetic pattern, in line with previous studies of intestinal metamorphosis in this species (Sterling *et al*., 2012) as well as stomach development in other anurans (Rovira *et al*., 1995). We additionally report that the larval stomach lining of *X. tropicalis* increases in complexity after onset of larval feeding. However, the stomach lining remains homogeneous throughout its anterior to posterior extent despite developing more numerous tubular glands, and the epithelium develops ciliated cells facing the lumen with numerous folds, gastric pits, and an apical mucosal lining. Feeding ceases during metamorphic remodeling, and by the end of metamorphosis the stomach of *X. tropicalis* is equipped with both fundic and pyloric regions suited to digest a carnivorous diet.

In *L. laevis*, the metamorphic transition from larva to adult is, with respect to many morphological features, typical for biphasic anurans—the larval tail is lost, limbs form, etc. However, features related to feeding diverge from this path, including the cranial skeleton and musculature, hyobranchial apparatus and gastrointestinal tract (Fabrezi & Lobo, 2009; Ziermann, *et al*., 2011; Fabrezi & Cruz, 2020). The larval gastrointestinal lining is not shed and instead remains functional during metamorphosis, corresponding to the larva’s ability to feed throughout this transition. However, between the onset of larval feeding soon after hatching and the completion of metamorphosis, the stomach increases in complexity in a manner similar to the increase in stomach complexity between early and late larval *X. tropicalis*. No new tissues form in *X. tropicalis* or *L. laevis* during the period of larval growth before metamorphosis; instead, existing tissues continue to increase in complexity–the epithelium and the glandular mucosa thicken and the glands increase in size and number. In the fundic region of *L. laevis*, the mucosa thickens and develops many more glands containing mucous neck cells, and the muscularis mucosa thickens to become visibly distinct from the serosa. In the pyloric region, the epithelium becomes more folded and the muscularis mucosa thickens. Despite not shedding and replacing its lining, the stomach of *L. laevis* does undergo slight morphological change during metamorphosis. The fundic region changes the most, though still relatively little compared to *X. tropicalis*, by the addition of the muscularis mucosa and an aglandular submucosa. The pyloric region only differs in the addition of the muscularis mucosa between the lamina propria and submucosa. This ongoing maturation of stomach microanatomy does not appear to interfere with feeding, as larvae feed continuously during larval growth and metamorphosis. This increase in stomach complexity also corresponds to simultaneous trends in increased pepsin production (Carroll *et al*., 1991).

The ability of *L. laevis* to feed continually throughout metamorphosis is enabled by its development of an adult stomach during embryonic development. However, the stomach is not entirely static after hatching. It retains the ancestral pattern of an increase in complexity during the larval period and incurs additional slight changes during metamorphosis.

### *Evolutionary significance of larval carnivory in* Lepidobatrachus laevis

Stomach development in *L. laevis* follows a unique ontogenetic trajectory that spans both the embryonic period and post-hatching metamorphosis. This novel gastric ontogeny allows *L. laevis* to have a megalophagous, carnivorous feeding strategy across both larval and adult phases. The consequent departure from the typical herbivorous feeding mode of larval anurans narrows the ecomorphological distance between larva and adult and may significantly reduce dietary partitioning between these two discrete life-history stages (Fabrezi, 2006, 2011; Fabrezi & Quinzio, 2008). *Lepidobatrachus laevis* is adapted to an extremely arid desert region, the Gran Chaco, which spans adjacent portions of Paraguay, Bolivia and Argentina. Adults occupy ephemeral pools formed during the rainy season but retreat into underground burrows and envelop themselves in epidermal cocoons to survive the dry season (Budgett, 1899; McClanahan, 1976; Ruibal & Thomas, 1988; Faivovich 2014). This life history is thought to impose significant selection pressure for tadpoles to shorten the larval period and quickly metamorphose into adults capable of surviving the annual period of estivation. Such pressure to quickly attain the adult stage likely outweighs the ancestral benefits of niche partitioning between larval and adult stages and underlies evolution of the larval carnivorous diet (Fabrezi, 2006, 2011; Fabrezi & Quinzio, 2008). Although carnivorous tadpoles are not unique to *L. laevis*, the species and its congeners are unique in the extremes to which they take larval carnivory. The stomach, for example, is capable of distending greatly, allowing tadpoles to eat prey nearly their own size. We have observed *Lepidobatrachus laevis* tadpoles to consume other tadpoles so large that the prey’s tail partially extends from heir mouths until it can be digested enough to be swallowed entirely. The adult-like morphology of the larval stomach reflects this extreme as well; even relative to carnivorous tadpoles of other species, such as *Ceratophrys*, the larval stomach of *L. laevis* is most similar to that of adults (Bloom, 2013; Fabrezi & Cruz, 2020).

The complex life cycle of *Lepidobatrachus laevis* includes a discrete larval phase, although one that includes some adult traits, including the stomach, that do not form until metamorphosis in most other anuran species. These changes in ontogenetic timing of morphological development constitute an obvious example of heterochrony (Hanken, 1993; Fabrezi, 2011). This raises the question: What underlying developmental mechanisms have mediated the heterochronic shift in the timing of stomach development during the evolution of *L. laevis?* Have metamorphic stomach development modules simply shifted to embryonic development? A temporal shift in development would require that the metamorphic modules be integrated with larval ones that initially pattern the gut during embryogenesis. Teasing apart mechanisms of stomach evolution in *L. leavi*s may help determine which ones are necessary for building the essential components of a gut in general, as opposed to those that underlie the formation of particular larval gastric specializations, such as a filter-feeding herbivore vs. a bulk-feeding carnivore. Previous studies have implicated both increased thyroid hormone signaling and reduced retinoic acid signaling as possible mechanisms (Bloom *et al*., 2013). The histological timeline of stomach development defined by this study lays the groundwork for follow-up studies of the genetic mechanisms behind frog stomach development by identifying important time points of developmental change. *Lepidobatrachus laevis* can serve as an important model for studying the mechanisms that underlie heterochrony in generating evolutionary change.

## Acknowledgements

We would like to thank Prof. Nanette Nascone-Yoder at North Carolina State University for allowing us to use her lab’s *Lepidobatrachus laevis* and for her support. We’d also like to thank Dr. Brent Wyatt and Dr. Mike Dush from the Nsacone-Yoder lab for their assistance and support in rearing the *L. laevis* tadpoles. Thank you to Dr. Zachary Lewis for consultation on histological techniques. We’d also like to express immense gratitude to Matthew Gage, the Hanken Lab Manager, and Cory Hahn, the Hanken Lab Animal Technician, for all of their technical support and excellent care of our *Xenopus tropicalis*.

